# Methylmap: visualization of modified nucleotides for large cohort sizes

**DOI:** 10.1101/2022.11.28.518239

**Authors:** Elise Coopman, Marleen Van den Broeck, Tim De Poorter, Geert Joris, Dennis W Dickson, Mojca Strazisar, Rosa Rademakers, Wouter De Coster

## Abstract

Methylmap is a tool developed for visualization of modified nucleotide frequencies per position, especially for large numbers of samples. Various input possibilities are supported, including the standardized BAM/CRAM files containing MM and ML tags.

**Availability and implementation:** Methylmap is written in Python3 and available through PyPI and bioconda. The source code is released under MIT license and can be found at https://github.com/EliseCoopman/methylmap.

## 1 Introduction

In recent years, the study of epigenetics has become a crucial topic of interest for understanding of biological functions. Nucleotide modifications have physiological functions such as transcription silencing, genomic imprinting, X-chromosome inactivation, regulation of expression during germline and embryonic development and repression of transposons (Greenberg and Bourc’his 2019) and are known to be involved in various disorders including cancer, neurodevelopmental disorders, neurodegenerative and neurological diseases and autoimmune diseases (Portela and Esteller 2010). Chemical and enzymatic methods have been used to investigate nucleotide modifications, mainly cytosine methylation, the most extensively studied epigenetic modification, with short-read sequencing (Zhao, Song et al. 2020). Third-generation single-molecule long-read sequencing technologies such as Oxford Nanopore Technology (ONT) and PacBio Single-Molecule Real-Time (SMRT) sequencing have transformed our ability for modification detection by enabling phasing reads into parental haplotypes, allowing for analysis of allelTechnology (ONT) and PacBioTechnology (ONT) and PacBioe-specific modification information across large distances (Kelleher, Murphy et al. 2018) (Liu, Fang et al. 2019). Due to ongoing developments in the field of long-read sequencing technologies, they are now being applied in population-scale (epigenetic) sequencing projects (De Coster, Weissensteiner et al. 2021). Several tools for visualization of nucleotide modification patterns in one or a limited number of individuals are available (De Coster, Stovner et al. 2020) (Pryszcz and Novoa 2021) (Su, Gouil et al. 2021) (Cheetham, Kindlova et al. 2022), however, to our knowledge, there is no software available for large cohort sizes.

## 2 Materials and methods

We developed methylmap, a tool for visualization of nucleotide modification frequencies per position for a large number of individuals and/or haplotypes in heatmap format for a genomic region of interest. Methylmap supports several input possibilities such as the recently standardized BAM or CRAM files with MM and ML tags, generated by for example remora (ONT) or primrose (PacBio). Other input possibilities are tab separated files from the nanopolish methylation caller (Simpson, Workman et al. 2017) and an own overview tab separated table with modification frequencies. A possibility to order input samples by experimental group is present. Tabix (Li 2011) is used for fast and efficient retrieval of the region of interest of the input files, making the tool applicable for large numbers of samples. Methylmap depends on the pandas (McKinney 2010), numpy (Harris, Millman et al. 2020), plotly (Plotly Technologies Inc. 2015), modbam2bed (ONT) and the argparse Python module (Van Rossum and Drake 2009). Optionally, there is the presence of a gene or transcript annotation track, supported by a GTF or GFF input file. Output of methylmap is an heatmap visualization in dynamic HTML format and its overview table with modification frequencies as tab separated file. Our software is implemented in Python (Van Rossum and Drake 2009), installable via PyPi, and available through Bioconda (Gruning, Dale et al. 2018).

The data for the example in figure 1 is generated from 87 frontal cortex brain samples (Mayo Clinic Brain Bank) and sequenced using the ONT PromethION. Data was base called using guppy (ONT), aligned with minimap2 (Li 2018) and phased by longshot (Edge and Bansal 2019). Methylation was detected with nanopolish (Simpson, Workman et al. 2017). The data was further processed using scripts originally available with methplotlib (De Coster, Stovner et al. 2020) resulting in two haplotype files per sample, making it possible to visualize allele-specific modifications. Figure 1 shows the GNAS locus (genomic location chr20:58,839,718-58,911,192). GNAS has known imprinted regions that alternate haplotype (Plagge and Kelsey 2006), as is visible in the heatmap. An annotation track, extracted from a GENCODE (Frankish, Diekhans et al. 2021) GFF file, displays the gene-exon structure. Generating figure 1 takes <10s and generates a 4.60 Mb plot in HTML format.

**Figure 1:**
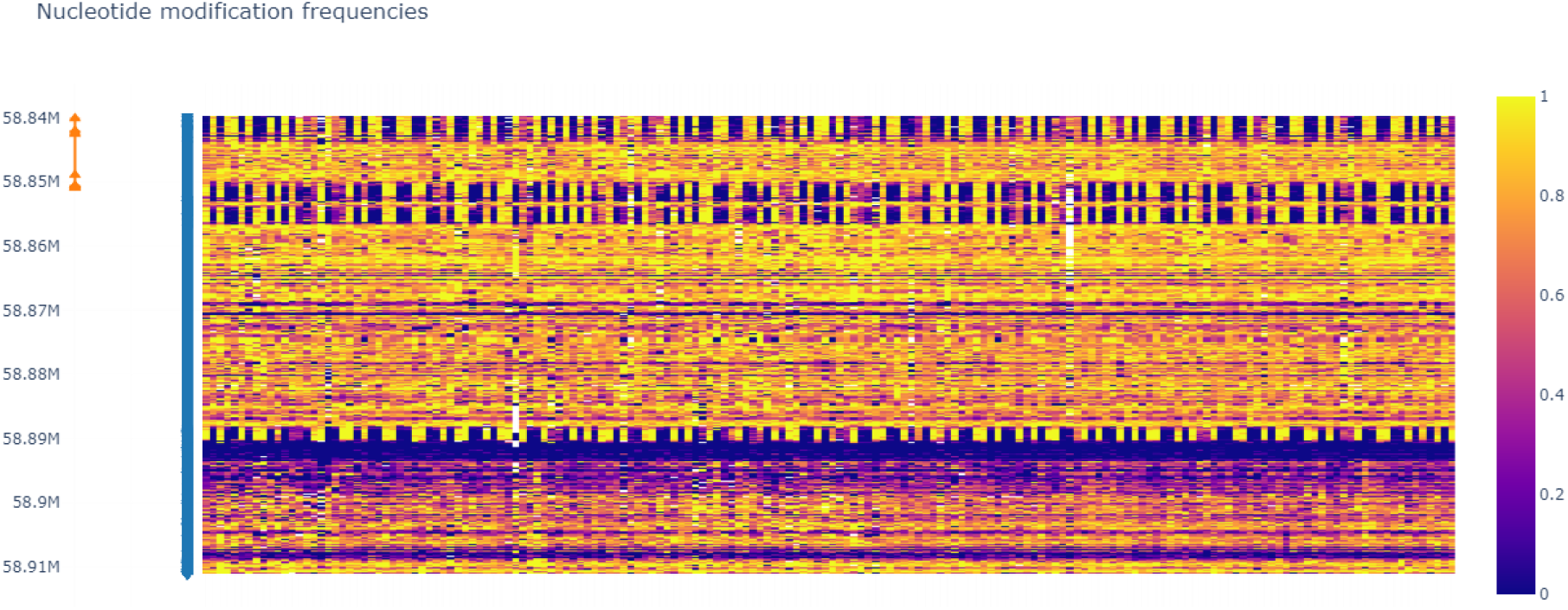
Methylation frequency of the known imprinted gene GNAS of 87 phased samples, showing the alternating haplotype imprinting pattern. At the left side, annotation of the region shows gene exon structure. The heatmap shows methylation frequencies with high methylated positions in yellow and low methylated positions in purple.

## 3 Conclusion

Over the past years, increased interest in epigenetic modifications resulted in extensive developments of technologies making it possible to perform population-scale epigenetic studies. We developed methylmap, a tool for visualization of modification frequencies of large cohort sizes. Methylmap is a technology-agnostic tool supporting standardized BAM/CRAM, nanopolish and own modification frequency table input, which can, together with other features, be expanded in the future.

## References

Cheetham, S. W., M. Kindlova and A. D. Ewing (2022). “Methylartist: tools for visualizing modified bases from nanopore sequence data.” Bioinformatics 38(11): 3109–3112.

De Coster, W., E. B. Stovner and M. Strazisar (2020). “Methplotlib: analysis of modified nucleotides from nanopore sequencing.” Bioinformatics 36(10): 3236–3238.

De Coster, W., M. H. Weissensteiner and F. J. Sedlazeck (2021). “Towards population-scale long-read sequencing.” Nat Rev Genet 22(9): 572–587.

Edge, P. and V. Bansal (2019). “Longshot enables accurate variant calling in diploid genomes from single-molecule long read sequencing.” Nat Commun 10(1): 4660.

Frankish, A., M. Diekhans, I. Jungreis, J. Lagarde, J. E. Loveland, J. M. Mudge, C. Sisu, J. C. Wright, J. Armstrong, I. Barnes, A. Berry, A. Bignell, C. Boix, S. Carbonell Sala, F. Cunningham, T. Di Domenico, S. Donaldson, I. T. Fiddes, C. Garcia Giron, J. M. Gonzalez, T. Grego, M. Hardy, T. Hourlier, K. L. Howe, T. Hunt, O. G. Izuogu, R. Johnson, F. J. Martin, L. Martinez, S. Mohanan, P. Muir, F. C. P. Navarro, A. Parker, B. Pei, F. Pozo, F. C. Riera, M. Ruffier, B. M. Schmitt, E. Stapleton, M. M. Suner, I. Sycheva, B. Uszczynska-Ratajczak, M. Y. Wolf, J. Xu, Y. T. Yang, A. Yates, D. Zerbino, Y. Zhang, J. S. Choudhary, M. Gerstein, R. Guigo, T. J. P. Hubbard, M. Kellis, B. Paten, M. L. Tress and P. Flicek (2021). “Gencode 2021.” Nucleic Acids Res 49(D1): D916–D923.

Greenberg, M. V. C. and D. Bourc’his (2019). “The diverse roles of DNA methylation in mammalian development and disease.” Nat Rev Mol Cell Biol 20(10): 590–607.

Gruning, B., R. Dale, A. Sjodin, B. A. Chapman, J. Rowe, C. H. Tomkins-Tinch, R. Valieris, J. Koster and T. Bioconda (2018). “Bioconda: sustainable and comprehensive software distribution for the life sciences.” Nat Methods 15(7): 475–476.

Harris, C. R., K. J. Millman, S. J. van der Walt, R. Gommers, P. Virtanen, D. Cournapeau, E. Wieser, J. Taylor, S. Berg, N. J. Smith, R. Kern, M. Picus, S. Hoyer, M. H. van Kerkwijk, M. Brett, A. Haldane, J. F. Del Rio, M. Wiebe, P. Peterson, P. Gerard-Marchant, K. Sheppard, T. Reddy, W. Weckesser, H. Abbasi, C. Gohlke and T. E. Oliphant (2020). “Array programming with NumPy.” Nature 585(7825): 357–362.

Kelleher, P., J. Murphy, J. Mahony and D. van Sinderen (2018). “Identification of DNA Base Modifications by Means of Pacific Biosciences RS Sequencing Technology.” Methods Mol Biol 1681: 127–137.

Li, H. (2011). “Tabix: fast retrieval of sequence features from generic TAB-delimited files.” Bioinformatics 27(5): 718–719.

Li, H. (2018). “Minimap2: pairwise alignment for nucleotide sequences.” Bioinformatics 34(18): 3094–3100.

Liu, Q., L. Fang, G. Yu, D. Wang, C. L. Xiao and K. Wang (2019). “Detection of DNA base modifications by deep recurrent neural network on Oxford Nanopore sequencing data.” Nat Commun 10(1): 2449.

McKinney, W. (2010). Data Structures for Statistical Computing in Python. Proceedings of the 9th Python in Science Conference. S. van der Walt and J. Millman. 445: 56–61.

Plagge, A. and G. Kelsey (2006). “Imprinting the Gnas locus.” Cytogenet Genome Res 113(1-4): 178–187.

Plotly Technologies Inc. (2015). “Collaborative data science.” from https://plot.ly.

Portela, A. and M. Esteller (2010). “Epigenetic modifications and human disease.” Nat Biotechnol 28(10): 1057–1068.

Pryszcz, L. P. and E. M. Novoa (2021). “ModPhred: an integrative toolkit for the analysis and storage of nanopore sequencing DNA and RNA modification data.” Bioinformatics.

Simpson, J. T., R. E. Workman, P. C. Zuzarte, M. David, L. J. Dursi and W. Timp (2017). “Detecting DNA cytosine methylation using nanopore sequencing.” Nat Methods 14(4): 407–410.

Su, S., Q. Gouil, M. E. Blewitt, D. Cook, P. F. Hickey and M. E. Ritchie (2021). “NanoMethViz: An R/Bioconductor package for visualizing long-read methylation data.” PLoS Comput Biol 17(10): e1009524.

Van Rossum, G. and F. L. Drake (2009). Python 3 Reference Manual, CreateSpace.

Zhao, L. Y., J. Song, Y. Liu, C. X. Song and C. Yi (2020). “Mapping the epigenetic modifications of DNA and RNA.” Protein Cell 11(11): 792–808.

